# Engineering of critical enzymes and pathways for improved triterpenoid biosynthesis in yeast

**DOI:** 10.1101/2020.04.03.023150

**Authors:** Hao Guo, Huiyan Wang, Yi-xin Huo

## Abstract

Triterpenoids represent a diverse group of phytochemicals, widely distributed in the plant kingdom with many biological activities. Recently, the heterologous production of triterpenoids in *Saccharomyces cerevisiae* has been successfully implemented by introducing various triterpenoids biosynthetic pathways. By engineering related enzymes as well as yeast metabolism, the yield of various triterpenoids is significantly improved from milligram-scale per liter to gram-scale level per liter. This achievement demonstrates that engineering of critical enzymes is considered as a potential strategy to overcome the main hurdles of translation of these potent natural products into industry. Here, we review strategies for designing enzymes to improve the yield of triterpenoids in *S. cerevisiae*, which is mainly separated into three aspects: 1. elevating the supply of the precursor—2,3-oxidosqualene, 2. optimizing triterpenoid-involved reactions, 3. lowering the competition of the native sterol pathway. And then we provide challenges and prospects on further enhancing the triterpenoid production in *S. cerevisiae*.

## Introduction

Triterpenoids are a subset of terpenoids that are widely distributed in plants, microbes, and marine organisms with various structures and applications reported to date [1], e.g. artemisinin, taxol, lycopene, stevioside, perilla alcohol, ginsenosides, and azadirachtin A (Figure 1). Terpenoids are classified into 7 subgroups: hemi- (C5), mono- (C10), sesqui- (C15), di- (C20), tri- (C30), tetra (C40) and poly- (C>40) terpenoids according to the number of isoprene units [2]. Triterpenoids consist of six isoprene units which are mainly synthesized from—2,3-oxidosqualene, except for a minority of triterpenoids generated from squalene and lanosterol. Most triterpenoids belong to secondary plant metabolites, which have been applied in the food, cosmetics, and pharmaceutical industry [3]. Recently, several triterpenoids have gained more attention, as they are the main active ingredients of many traditional medicines which contain a variety of pharmacological compounds, such as ginsenosides, betulinic acid, and oleanolic acid [4]. For example, ginsenosides are the main bioactive component in Asian and American ginseng that benefits the central nervous, cardiovascular, endocrine, and immune systems [5, 6]. Besides, betulinic acid is a very promising anti-tumor agent which can suppress tumor growth and induce apoptosis in human melanoma cells and other cancer cells, with relatively low cytotoxicity against healthy cells [7].

**Figure 1.**
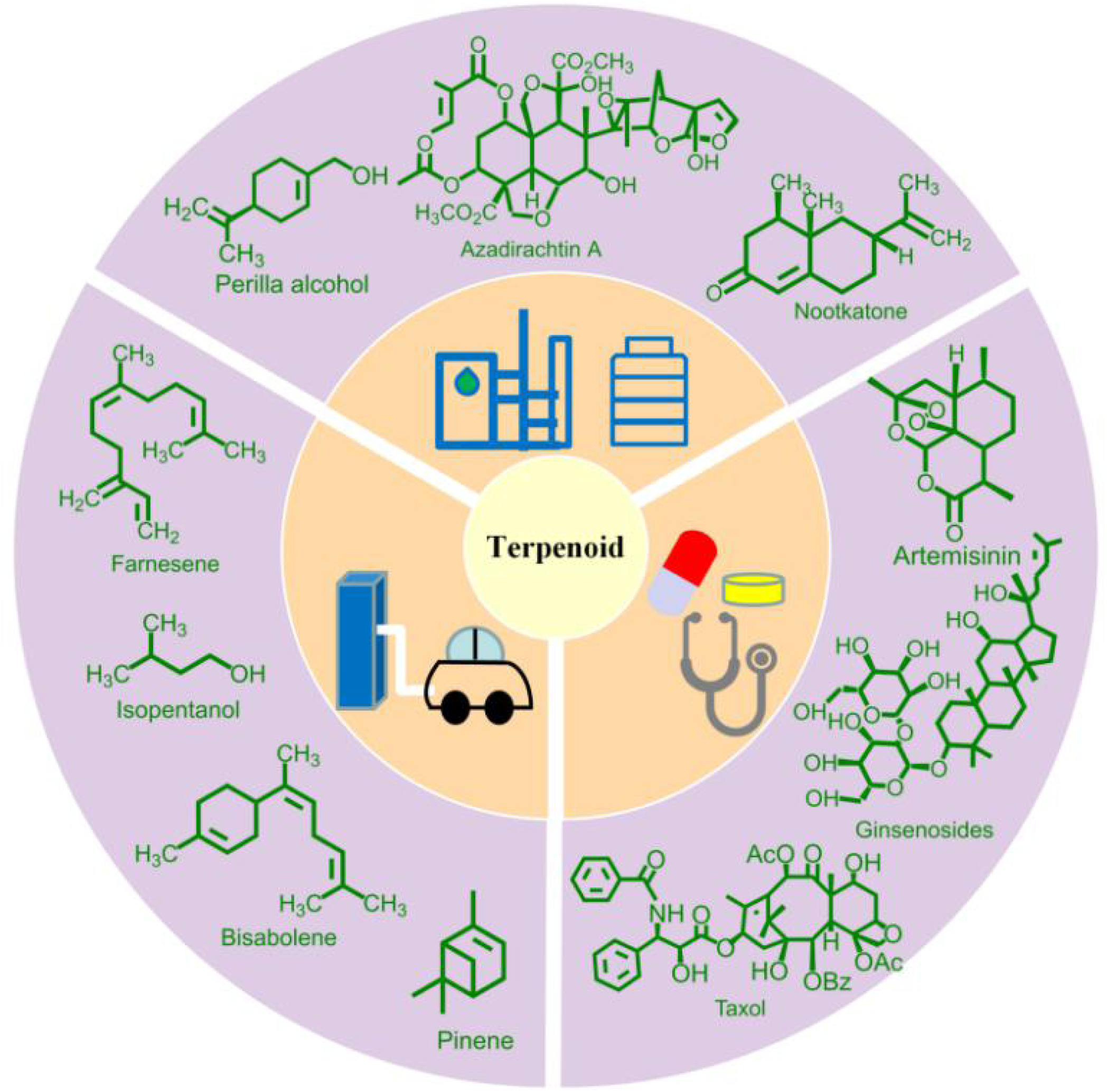
Structures and application of terpenoids in nature.

Typically, triterpenoids are produced by extraction from plant materials. However, the supply of triterpenoids is considerably limited by slow-growing plants and low extraction efficiency. Chemical synthesis is an infeasible way to substitute the plant extraction due to the complex skeletal characteristics and chiral centers of these molecules. For instance, azadirachtin A is a complex tetranor-triterpenoid from the Neem (*Azadirachta indica*), which is regarded as active ingredients of many pesticides [8]. The total chemical synthesis of azadirachtin A was completed after 22 years, which was required for 71 steps of reactions. Nonetheless, the final yield was only 0.00015% [9]. Hence, the microbial synthesis is an attractive alternative for the production of triterpenoids. Compared with current extraction from plant sources, this strategy possesses several advantages, such as short-cycle manufacture, land-saving, and controllable culture conditions.

Up to now, the production of triterpenoids has been successfully implemented in various microbes, such as *E. coli, S. cerevisiae, Y. lipolytica, P. pastoris. S. cerevisiae* is considered as an efficient homologous recombination system that can provide not only sufficient 2,3-oxidosqualene through its native sterol pathway but also the endoplasmic reticulum (ER) membrane for expression of appropriate enzymes. Hence, *S. cerevisiae* is predominately utilized in the heterologous production of triterpenoids. With the application of various metabolic engineering approaches on *S. cerevisiae*, the yield of triterpenoids, especially for dammarene-type, has been increased to the 10 gram-scale level per liter. However, the production of other types of triterpenoids is relatively lower, such as lupane-type, oleanane-type, and ursane-type triterpenoids. To sum up, there is still a large gap to realize the industrial production in place of extraction from plant sources. Here, we review the current progress in the elucidation of strategies to increase the production of triterpenoids as well as the modification of triterpenoid-related enzymes, and then we discuss challenges and perspectives on further enhancing the triterpenoids biosynthesis in *S. cerevisiae*

## Triterpenoid biosynthetic pathway

### 2,3-Oxidosqualene biosynthesis

The triterpenoid biosynthetic pathway can be mainly separated into two segments. 1. biosynthesis of the linear C30 molecule-2,3-oxidosqualene (Figure 2). 2. the formation of cyclic triterpenes based on the 2,3-oxidosqualene and further modification by other enzymes (Figure 3). Hence, 2,3-oxidosqualene is considered as a branch-point precursor in the synthesis of the linear and cyclic compounds during the triterpenoid biosynthesis. In general, there are two routes involved in the biosynthesis of 2,3-oxidosqualene—the mevalonate (MVA) pathway and methylerythritol-phosphate (MEP) pathway. However, in *S. cerevisiae*, the MVA pathway starting from the acetyl-CoA is the unique way to synthesize 2,3-oxidosqualene. Isopentenyl diphosphate (IPP) and dimethylallyl pyrophosphate (DMAPP) are end-products of the MVA pathway that are two fundamental building blocks for 2,3-oxidosqualene biosynthesis. Two molecules of IPP and one molecule of DMAPP are condensed to generate farnesyl diphosphate (FPP) by farnesyl diphosphate synthase, and two FPP molecules are further merged by squalene synthase to produce squalene that is oxidized to 2,3-oxidosqualene by squalene epoxidase. In *S. cerevisiae*, lanosterol is the only product of the cyclization of 2,3-oxidosqualene, which is an essential precursor for the sterol biosynthesis.

**Figure 2.**
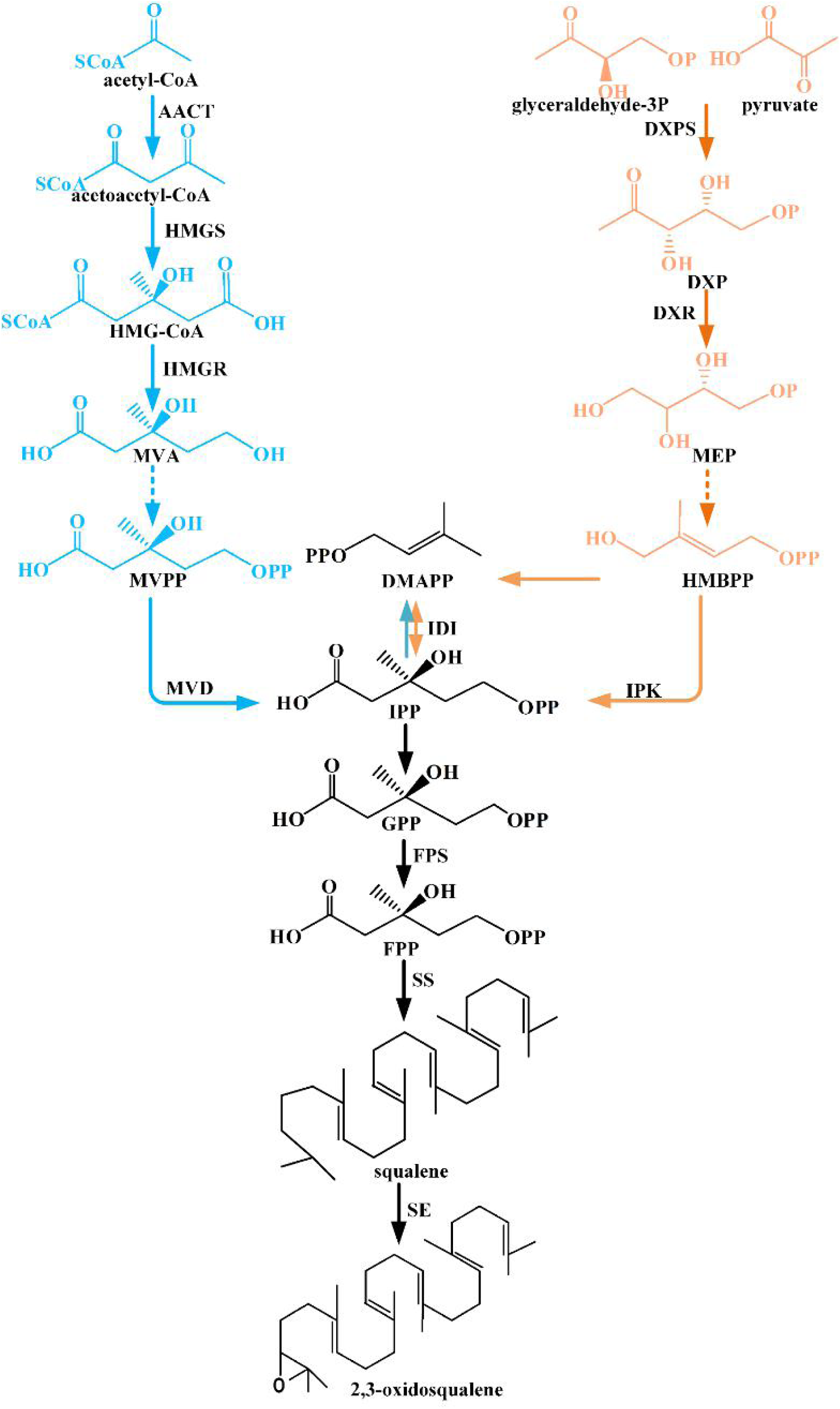
2,3-Oxidosqualene biosynthetic pathway. AACT, acetyl-CoA C-acetyltransferase; HMGS, 3-hydroxy-3-methylglutaryl-CoA synthase; HMGR, 3-hydroxy-3-methylglutaryl-CoA reductase; MVD, mevalonate diphosphate decarboxylase; DXPS, 1-deoxy-D-xylulose-5-phosphate synthase; DXR, 1-deoxy-D-xylulose-5-phosphate reductoisomerase; IPK, 4-hydroxy-3-methylbut-2-en-1-yl diphosphate reductase; IDI, isopentenyl-diphosphate delta-isomerase; FPS, farnesyl diphosphate synthase; SS, squalene synthase; SE, squalene epoxidase; HMGCoA, 3-hydroxy-3-methylglutaryl-CoA; MVA, mevalonate; MVPP, diphosphomevalonate; DXP,1-deoxy-D-xylulose-5-phosphate; MEP, methylerythritol phosphate; HMBPP, 1-Hydroxy-2-methyl-2-butenyl 4-diphosphate; IPP, isopentenyl diphosphate; GPP, geranyl diphosphate; FPP, farsenyldiphosphate; DMAPP, dimethylally diphosphate.

**Figure 3.**
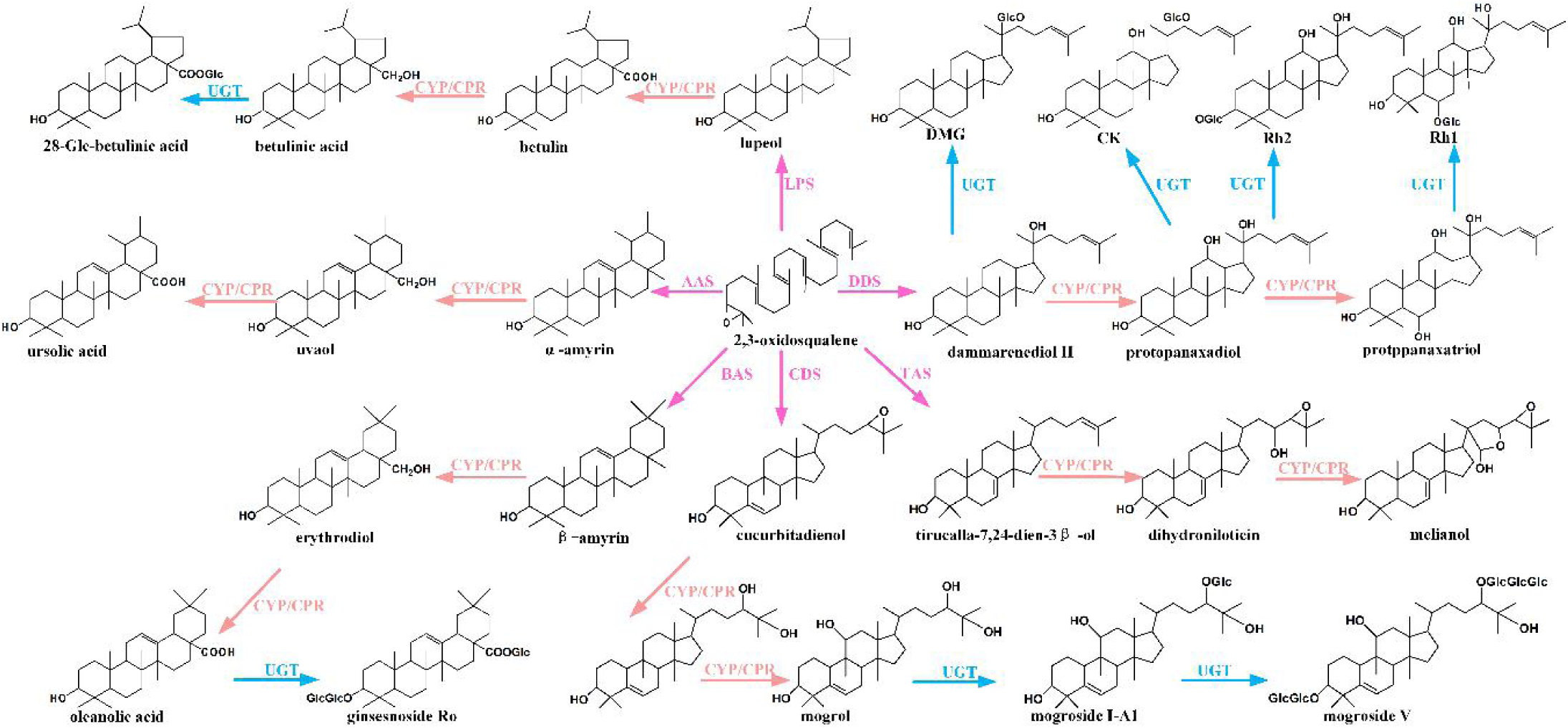
Triterpenoid saponins biosynthetic pathway. LPS, lupeol synthase; AAS, α-amyrin synthase; DDS, dammarenediol II synthase; BAS, β-amyrin synthase; CDS, cucurbitadienol synthase; TAS, tirucalla-7,24-3β-ol synthase. CYP, cytochrome P450; CPR, cytochrome P450 reductase; UGT, UDP-glycosyltransferase.

### 2,3-Oxidosqualene cyclases

In plants, the cyclization of 2,3-oxidosqualene is the first committed step for triterpenoid biosynthesis which is catalyzed by a variety of oxidosqualene cyclases (OSCs). To date, numerous OSCs have been identified in plants and heterologously expressed in engineered *E. coli* and 2,3-oxidosqualene producing yeast strains (Table 1). β-Amyrin, and lupeol synthases are the most abundant enzymes in the plant kingdom. In contrast, α-amyrin, dammarenediol II, thalianol, and taraxerol synthases are uncommon, especially as dammarenediol II synthase is only discovered in *Panax ginseng* and *Panax notoginseng*. These enzymes share the common conserved regions and the similar sequence. By site-specific mutagenesis of key amino acids in conserved regions, the function of these enzymes can be even switched to some extent. For example, L256W mutation of lupeol synthase from *Olea europaea* can result in the formation of β-amyrin rather than lupeol, whereas W259L mutation of β-amyrin synthase from *Panax ginseng* produces β-amyrin instead of lupeol [10](Figure 4).

**Table 1.**
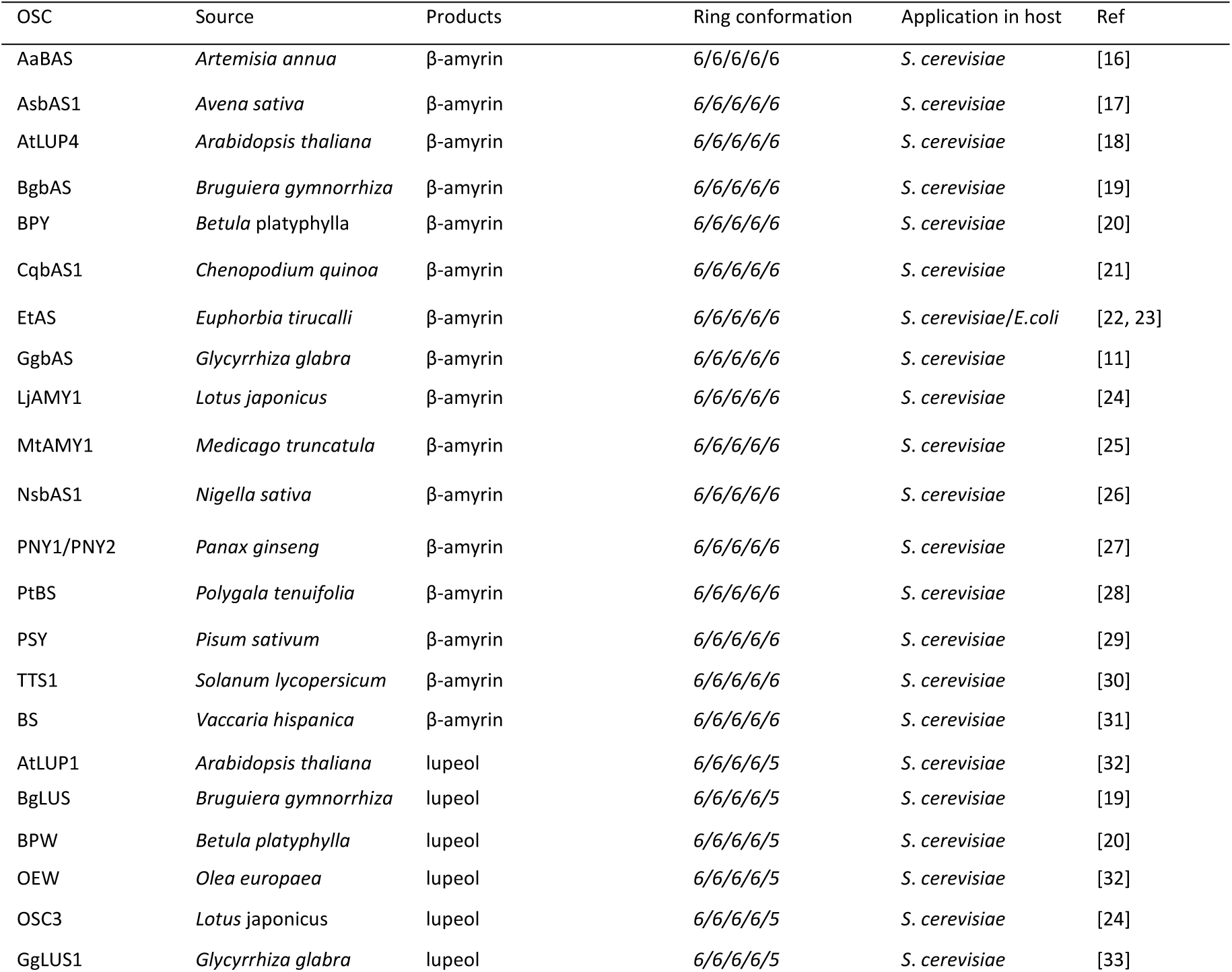
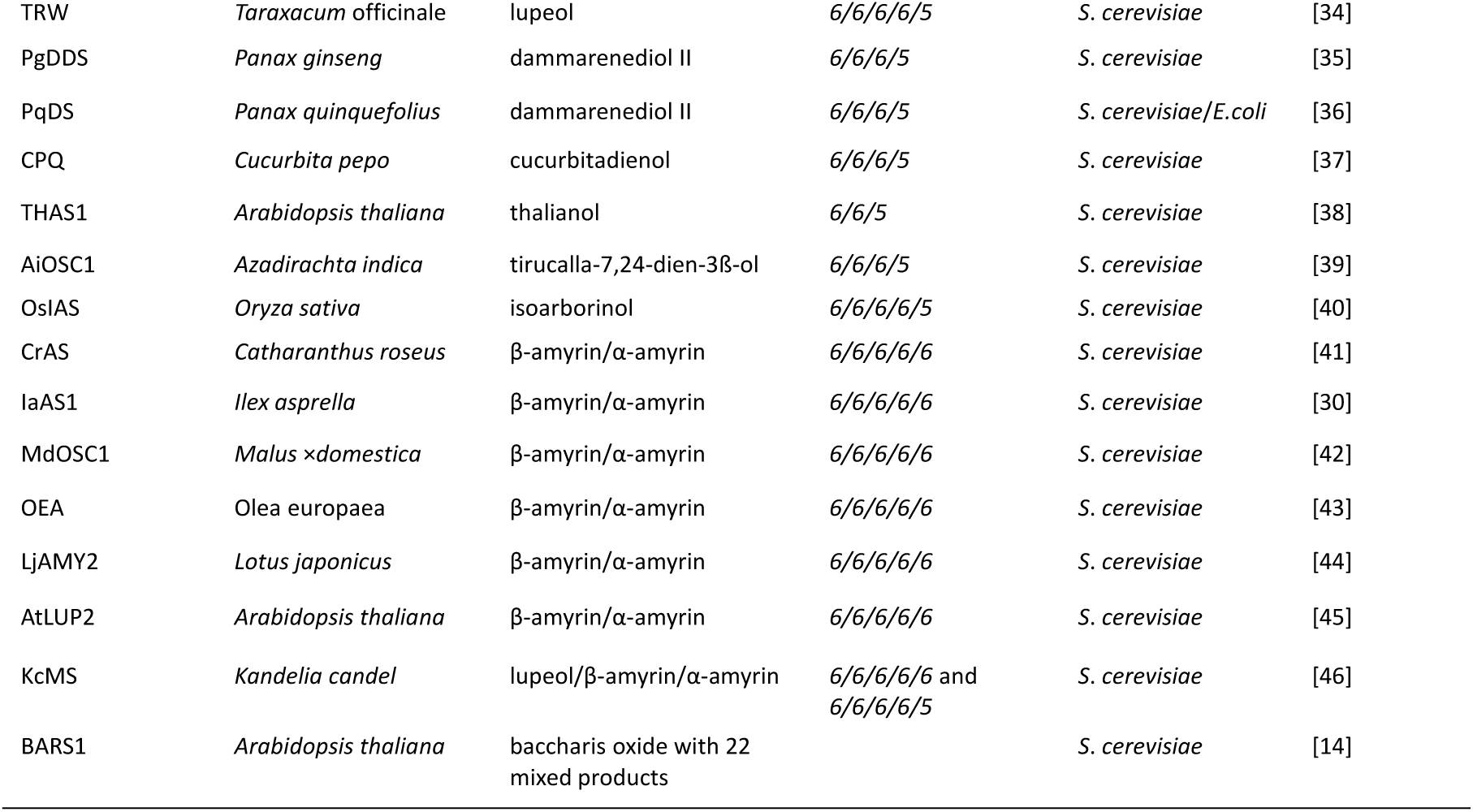
Verified 2,3-oxidosqualene cyclases for triterpene synthesis

**Figure 4.**
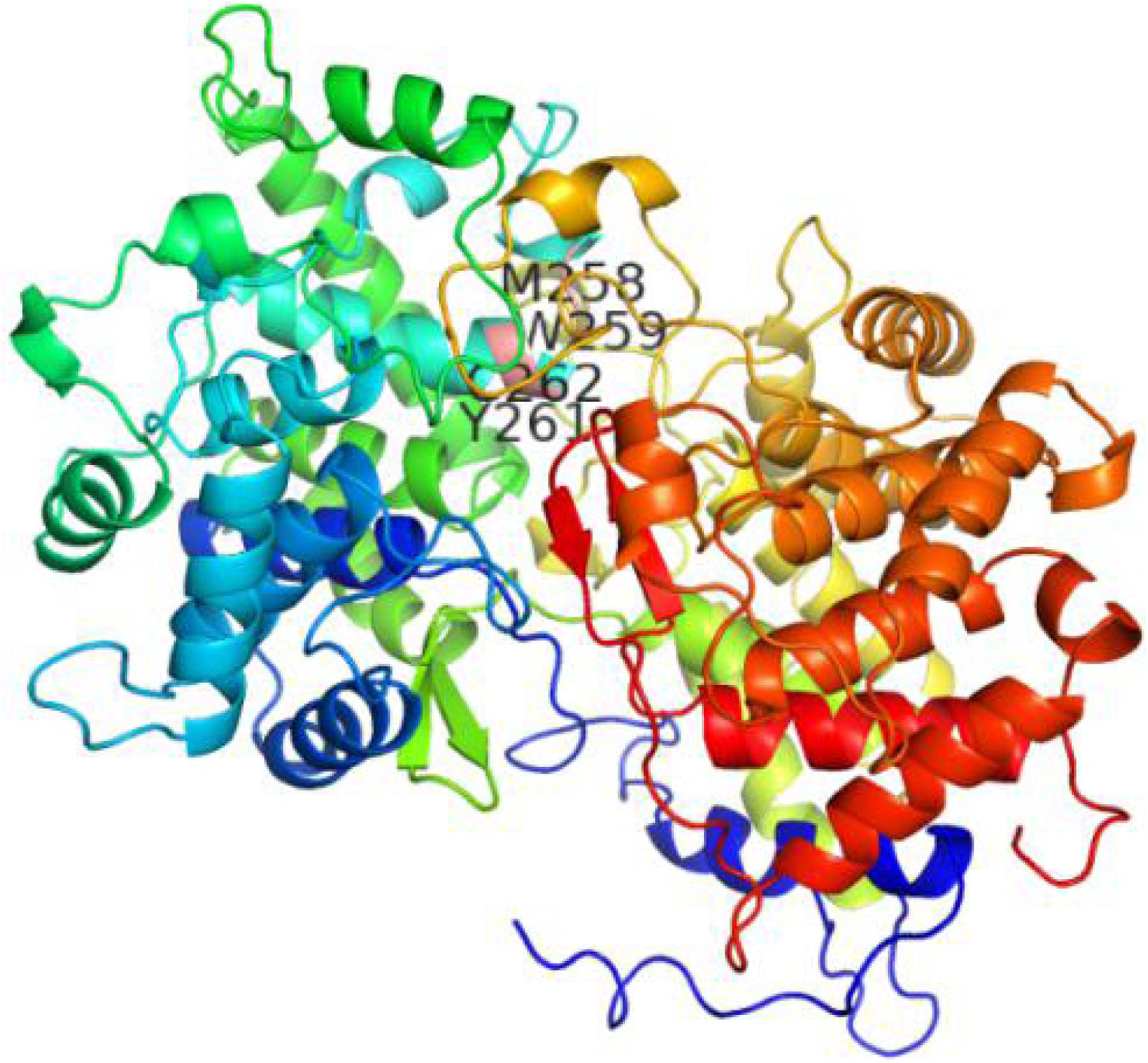
Site-specific mutagenesis of key amino acids in conserved regions of β-amyrin synthase (PNY).

The heterologous production of triterpenes has been achieved by the expression of several OSCs in *S. cerevisiae*. However, the yield of triterpenes varies depending on different OSCs of plant origin. For instance, the β-amyrin synthase from *G. glabra* yields more β-amyrin compared with other species, e.g. *A. annua, P. sativum, G. glabra*, and *P. ginseng* [11]. Also, α-amyrin synthase from *Malus* × *Domestica* has been characterized to exhibit high levels of specific activity which led to 5-fold increase on the yield of α-amyrin [12]. Hence, numerous isozymes provide multiple choices for metabolic engineering which is beneficial for triterpenoid production in *S. cerevisiae*.

Also, some OSCs are identified as multifunctional enzymes with mixed end-products [13]. In particular, baccharis oxide synthase from *A. thaliana* can make baruol as the primary product with up to 22 additional by-products through the cyclization of 2,3-oxidosqualene [14]. However, the mechanism of generation of mixed-products is unclear, it is probably due to the deprotonation of numerous sites during the cyclization process [15]. The metabolic diversity may be caused by the evolution, which allow plants to produce various triterpenoids against disease or insect attack. Nevertheless, it seriously influences the concentration of targeted products and causes a complicated purification process.

### P450 monooxygenases

P450 monooxygenases decorate basic triterpene skeletons by the introduction of hydroxyl, ketone, aldehyde, carboxyl, or epoxy groups. To date, a large number of P450s have been characterized as triterpene-oxidizing enzymes in various plant species (Table 2). Based on homology and phylogenetic criteria, the P450 family has been classified into 10 clans: CYP71, CYP72, CYP85, CYP86, CYP51, CYP74, CYP97, CYP710, CYP711, and CYP727 [47]. Many P450s exhibit substrate diversity compared with OSCs. Typically, pentacyclic triterpenoids, α-amyrin, β-amyrin, and lupeol, can be oxidized at the position of C12, C13, C-24, C-28, C-30, to generate mixed corresponding products by various P450s, such as CYP51H10, CYP93E1, CYP72A63, CYP716A12, and CYP716A15. A subgroup of the CYP85 clade, the CYP716, dominates the oxidation of triterpene skeletons in plants [48]. In contrast, a tetracyclic triterpenoid—dammarenediol II can be merely oxidized at the C-12 and C-6 position in sequence, to generate protopanaxadiol and protopanaxatriol respectively catalyzed by CYP716A47 and CYP716A53v2 [49].

**Table 2.**
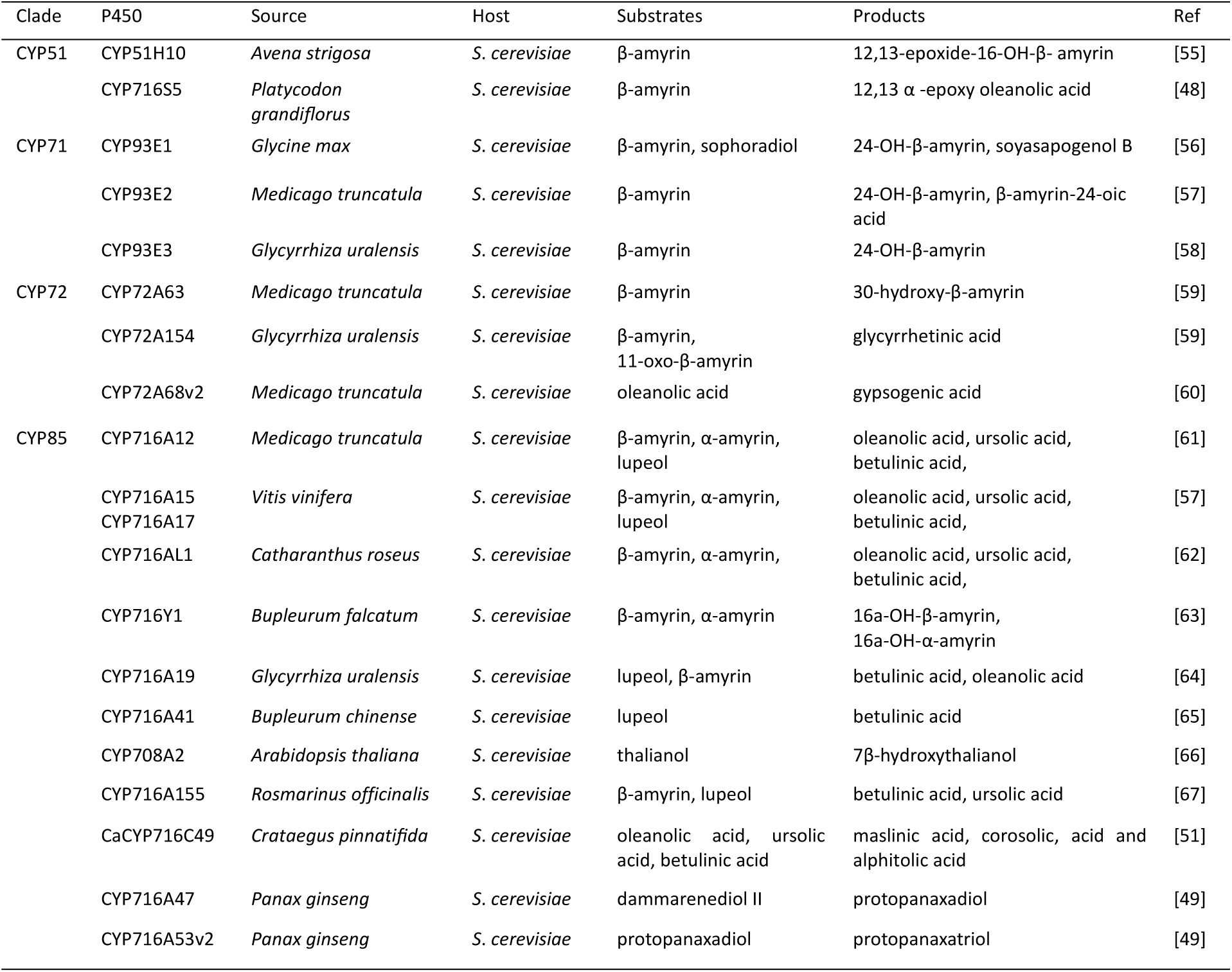
Plant triterpenoid-related cytochrome P450s in yeast

P450 typically catalyze the following reaction: RH + O_2_ + 2e^−^ + 2H^+^ → ROH + H_2_O. This reaction relies on the P450 reductase (CPR) for electron transfer as well as NAD(P)H for the electron donor. In some cases, cytochrome b_5_ is required as an alternative electron donor for P450-catalyzed reactions [50]. The formation of mixed end-products is possibly due to P450-CPR pairs, the lack of appropriate cytochrome b_5_, and the insufficiency of NADPH and oxygen supply.

As the numerous P450s and CPRs have been characterized in plants, the varying P450-CPR pairs were examined for improving the yield of triterpenoids. Dai et al. introduced different combinations of CYP450s and CPRs in yeast, such as MtCYP716A12 from *M. truncatula*, CrCYP716AL1 from *C. roseus*, VvCYP716A15 from *V. vinifera*, CPR from *A. thaliana, X. dendrorhous, L. japonicas, P. ginseng*, and *V. vinifera*. Finally, the yeast strain bearing the CYP716A15 and CPR from *V. vinifera* exhibited the highest yield of lupane-type triterpenoids [51]. In addition to screen different combinations of P450-CPR pairs, the protein engineering strategies have been utilized to modify the P450 reaction. For instance, a fusion protein strategy was applied on the protopanaxadiol synthase which was fused to its cofactor-truncated CPR1 from *A. thaliana* (ATR1, 46 amino acids were deleted at the N-terminus). The fusion enzyme (PPDS-ATR1) leads to a 71% increase in protopanaxadiol production compared with the separate expression of *PPDS* and *ATR1* genes [52]. Besides, an additional cytochrome b_5_ might benefit or inhibit the reaction depending on different P450s [53]. For example, CYP71AV1 is an amorphadiene oxidase which catalyzes three oxidation reactions from amorpha-4,11-diene to artemisinic acid, and expression of a cytochrome b_5_ from can benefit this reaction, which can increase the yield of artemisinic acid [54].

### UDP-glycosyltransferases

Triterpenoid saponins are generated by the glycosylation of triterpenoids, which is realized by transferring a sugar from UDP-sugar catalyzed by UGTs [68]. This modification step confers such compounds more water solubility, stability, and pharmacological properties. The group UGTs 71, 73, 74, 85, 91, and 94, are thought to be involved in the glycosylation of triterpenoid scaffolds (Table 3)[69]. Similar to P450s, UGTs also shows regioselectivity which catalyzes the glycosylation of triterpenoids at the position of C-3, C-28, C-6, C-20, and C-22 [70].

**Table 3.**
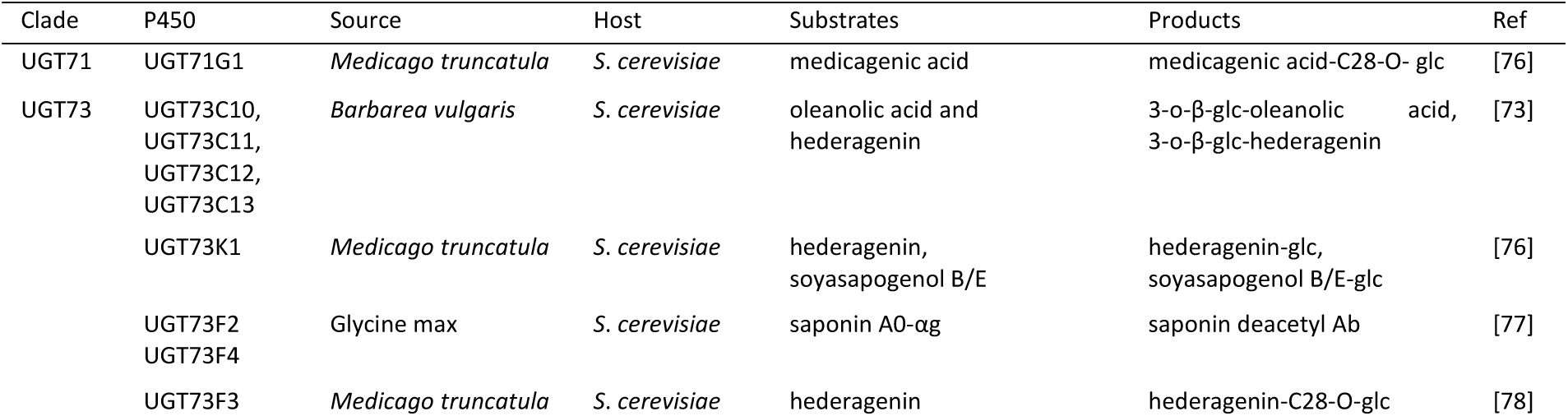
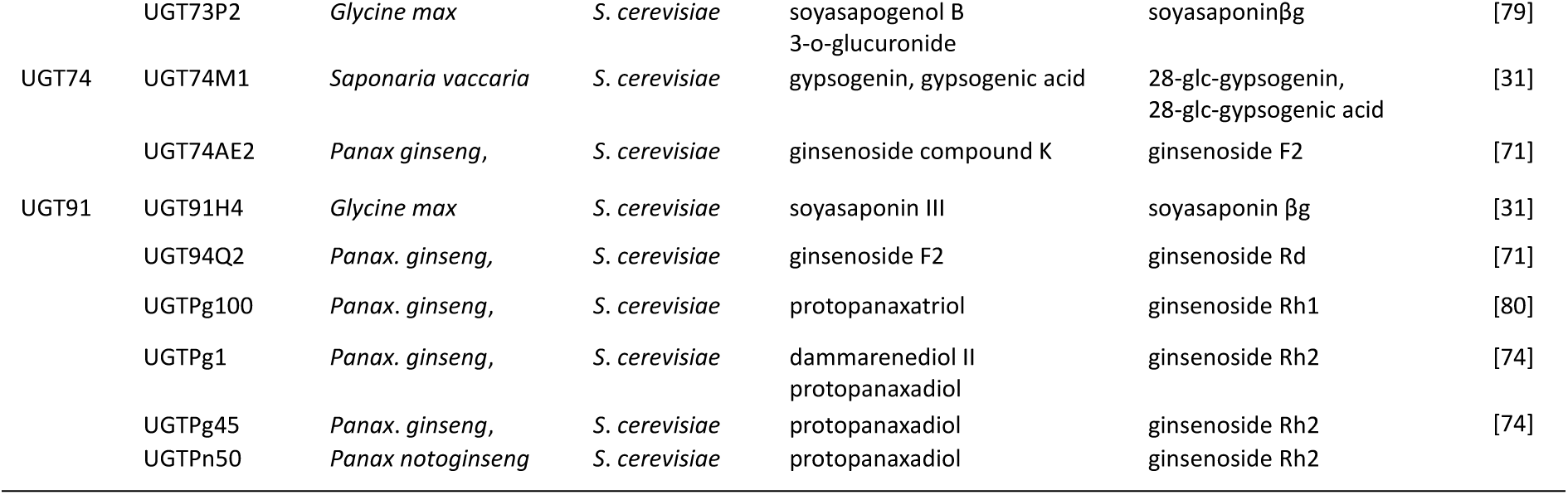
Plant UGTs verified in yeast

To date, numerous dammarene-type related UGTs have been identified in *P. ginseng*, which considerably enrich the structural diversity of ginsenosides. For example, PgUGT74AE2 can decorate the C3 hydroxyl groups of the protopanaxadiol and ginsenoside compound K to form Rh2 and F2 [71]. Moreover, the unnatural ginsenoside—3β,12β-Di-O-glc-protopanaxadiol has been synthesized from protopanaxadiol by UGT109A1 from *B. subtilis* [72]. In contrast, the pentacyclic triterpenoids-oleanolic acid and betulinic acid are mainly glycosylated by the UGT73 family (such as UGT73C10, UGT73C13) at their position of C-3 and C-28, to respectively generate 3-glc-oleanolic acid, 28-glc-betulinic acid [73].

Although the expression of UGTs has been successfully implemented in engineered yeast. the yield of these compounds is considerably low, only reaching to the milligram per liter. However, with the application of random mutations in the UGTPg45 gene, 2.25 g/L ginsenoside Rh2 in 10 L fed-batch fermentation was obtained based on the high-yield of its precursor in yeast [74]. Furthermore, the UGT73C5 coupled with sucrose synthase (from *A. thaliana*) was applied to generate the ginsenoside Rh2 from protopanaxadiol *in vitro*. This is due to that sucrose synthase can convert sucrose and UDP into UDPG which is indispensable for the glycosylation. Finally, the highest production of ginsenoside Rh2 reaches to 3.2 g/L by gradually adding protopanaxadiol in the reaction [75].

## Engineering yeast metabolism for improving the triterpenoids production

### Enhancing the precursor supply

A metabolic engineering approach is a powerful tool that can improve the yield and productivity of target products, reduce by-products, and increase cell viability in various microbes. Up to now, the most successful example is engineering *S. cerevisiae* for the heterologous production of artemisinic acid, which is the precursor of artemisinin–an effective drug for people suffering from malaria [81]. To achieve this, *S. cerevisiae* was engineered by up-regulating the expression of genes (overexpression of *tHMG1* gene) in the MVA pathway, repressing the competing reaction (transcriptional restriction of *ERG9* expression), improving the catalytic efficiency of P450 reactions (additional expression of *ADH1* and *ALDH1* genes from *A. annua*). Finally, 25 g/L artemisinic acid was achieved by an ethanol pulse feed process which has been applied in industrial production [54](Figure 5). This case sheds the light on the production of triterpenoids. Currently, the strategy for improving the production of triterpenoids is mainly implemented in three steps: 1. increasing the supply of the precursor—2,3-oxidosqualene. 2. optimizing the triterpenoid biosynthetic pathway. 3. lowering the competition of the native sterol pathway (Table 4).

**Table 4.**
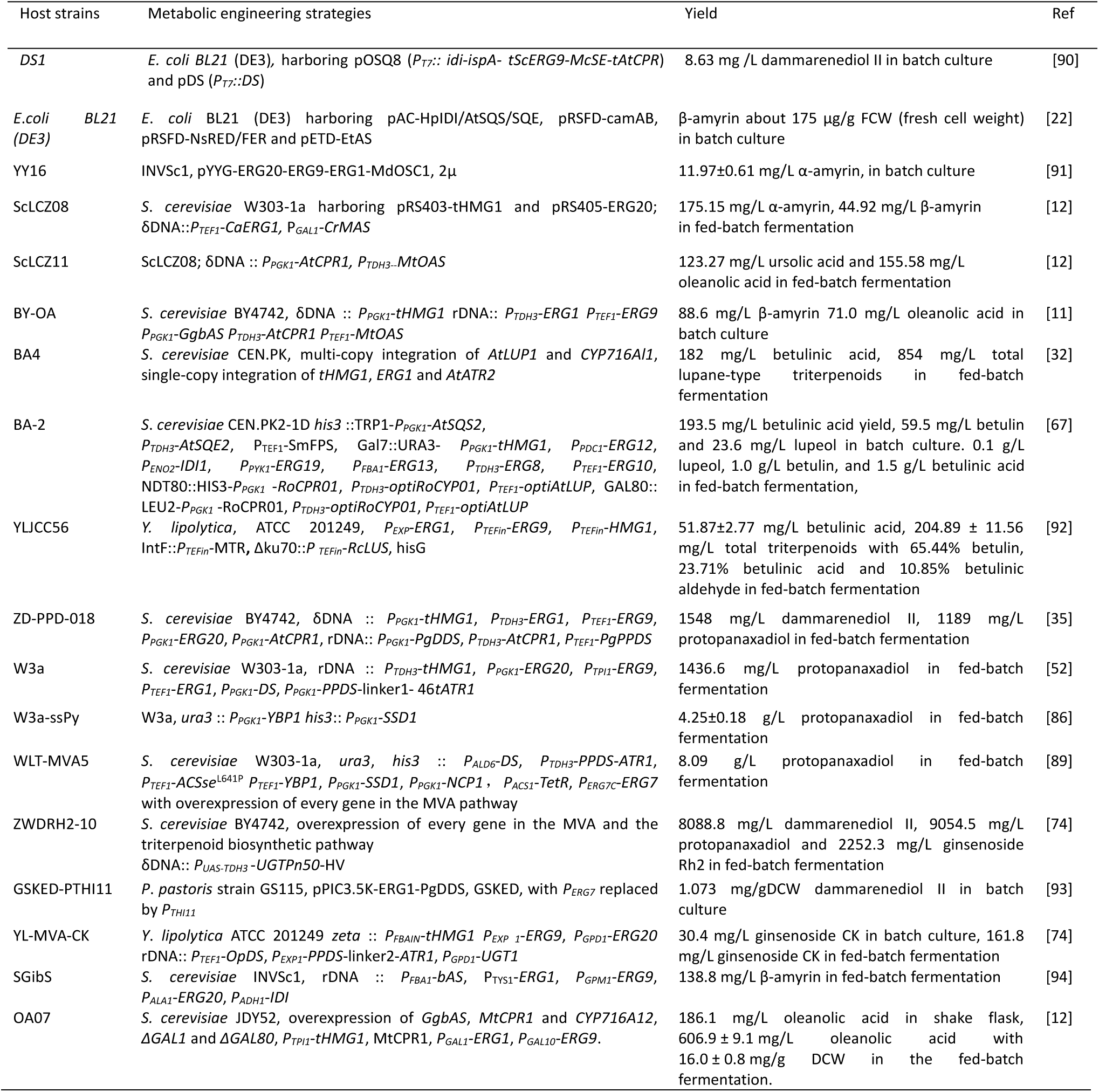
Metabolic engineering for production of triterpenoids in various microbes

**Figure 5.**
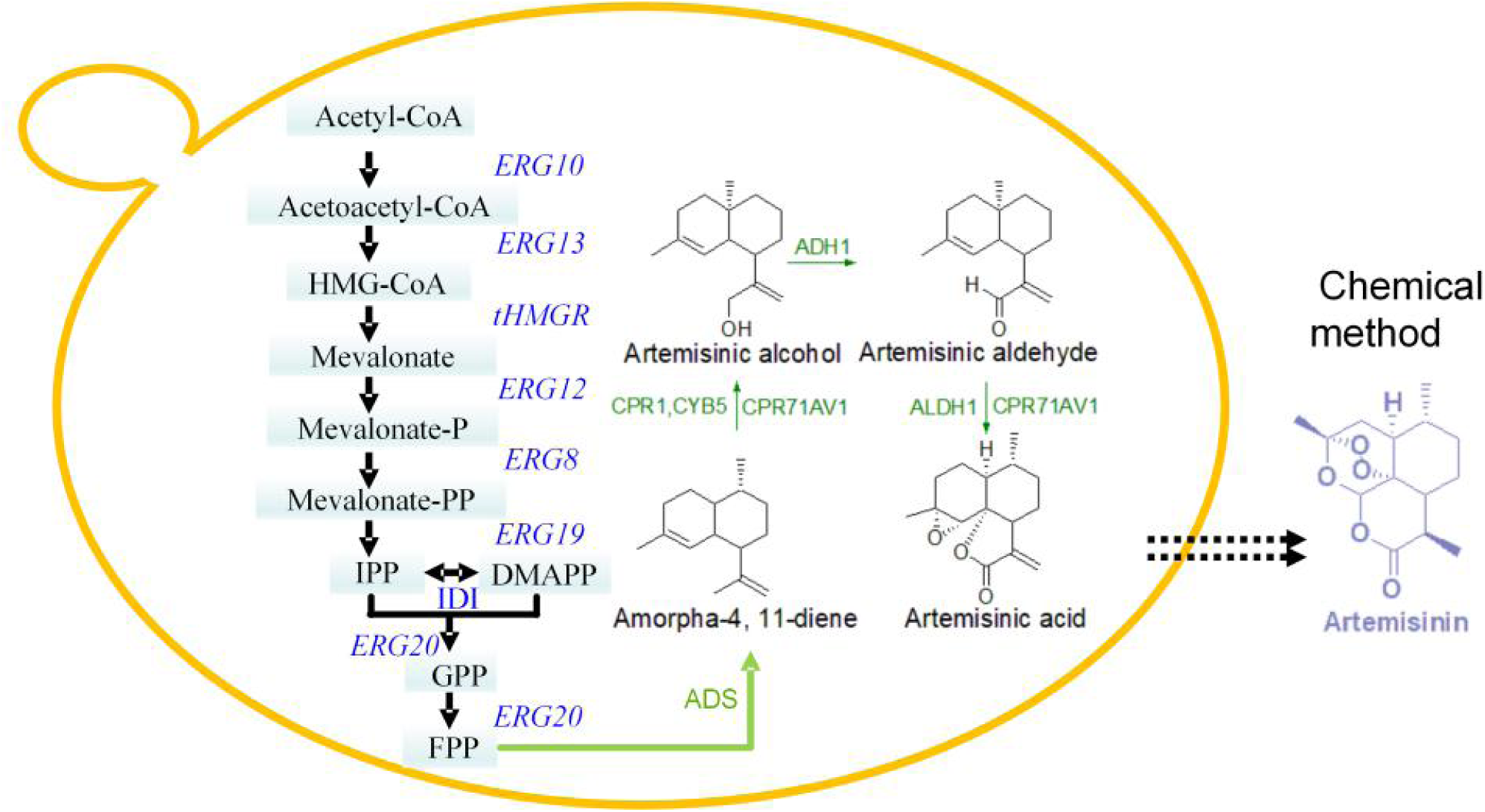
Overview of artemisinic acid production pathway in *S. cerevisiae*. The genes in the pathway for biosynthesis of FPP are shown in blue, genes encoding the full three-step oxidation of amorphadiene to artemisinic acid from *A. annua* are shown in green. DMAPP: dimethylallyl diphosphate; FPP: farnesyl diphosphate; IPP: isopentenyl diphosphate. tHMGR: truncated HMG-CoA reductase. CYP71AV1, CPR1 and CYB5 oxidize amorphadiene to artemisinic alcohol, aldehyde, and aldehyde. ADH1 oxidizes artemisinic alcohol to artemisinic aldehyde; ALDH1 oxidizes artemisinic aldehyde to artemisinic acid.

Overexpression of the *tHMG1* gene results in the accumulation of squalene, which is a common and efficient strategy for enhancing triterpenoid production [82, 83]. It is due to that the reduction of hydroxymethylglutaryl-CoA (HMG-CoA) to mevalonate catalyzed by HMG-CoA reductase is generally considered as a rate-limiting step in the MVA pathway. As squalene is an indirect precursor of triterpenoids biosynthesis. squalene epoxidase (encoded by *ERG1* gene) is also deemed to catalyze another rate-limiting step in the biosynthesis of triterpenoids. Dai et al. demonstrated that overexpression of the *ERG1* gene (encoding squalene synthase) could result in a 10-fold increase in the yield of protopanaxadiol [35].

Besides the rate-limiting step, elevating the expression level of all enzymes involved in the MVA pathway is also beneficial for increasing the carbon flux in yeast. Up to 18 g/L dammarene-type triterpenoids (dammarenediol II and protopanaxadiol) was achieved based on multi-copy integration of the whole 10 genes in the mevalonate pathway in the chromosome of *S. cerevisiae* using delta DNA site [74]. However, a similar strategy seems to have limited effects on the production of oleanane-type, ursane-type, and lupane-type triterpenoids. With the high expression level of all genes of the MVA pathway as well as CYP716A155 and CPR, approximately 1 g/L betulin and 1.5 g/L betulinic acid were achieved though the fed-batch fermentation [67]. Also, 600 mg/L oleanolic acid and 130 mg/L ursolic acid are respectively the highest yield achieved through the fed-batch fermentation reported to date [12], which is much lower than dammarene-type triterpenoids. It is probably caused by different OSCs, in which dammarenediol synthase exhibited higher enzyme activities than other cyclases in *S. cerevisiae*. Despite this, higher-level expression of rate-limiting enzymes, as well as other enzymes, is considered as an essential and efficient strategy for increasing the production of triterpenoids.

The reactions catalyzed by P450s and CPR are also vital for achieving the high-yield of triterpenoids, which requires extra carbon, NADPH and oxygen. The poor coupling of P450-CPR may result in the excess generation of the reactive oxygen species (ROS) which can damage DNA, proteins, lipids, even cause apoptosis in the cell [24]. Hence, regulation of yeast metabolism by engineering related enzymes is an alternative to benefit the yield of triterpenoids. Zhao et al. showed that expression of *SSD1* and *YBP1* genes (involved in the cellular response to oxidative stress), effectively decreased the ROS level and increased cell viability, which further improved the efficiency of protopanaxadiol biosynthesis [23]. Enhancing the supply of acetyl-CoA by overexpression of both the *ACS1* gene (acetyl-CoA synthetase) from *S. enterica* and the native *ALD6* gene could also increase the amorphadiene (a sesquiterpene) production [84]. Moreover, the deletion of *GDH1* gene (encoding glutamate dehydrogenase) enhanced the available NADPH in the cytosol which also resulted in an approximately 85% increase in the production of cubebol (a sesquiterpene) [85].

### Restraining the native sterol pathway

*S. cerevisiae* is not only used for triterpenoid but also for other terpenoid production based on intermediates of the MVA pathway. Hence, the competition between the native pathway and the heterologous pathway is always regarded as a hurdle for achieving the high yield of targeted products. The strategy for down-regulation of protein activity has been already applied to other metabolic nodes of the MVA pathway, such as Erg9p and Erg20p. Repressing the *ERG9* expression by replacing its native promoter with the copper-regulated promoter can efficiently redirect the carbon flow to the production of artemisinic acid [54]. Also, the ER-associated protein degradation system was applied to shorten the half-life of Erg9p which could improve the production of trans-nerolidol (a sesquiterpene) by 86% [86]. For the downregulation of *ERG20* gene, the expression of Erg20p was weakened by N-degron mediated protein degradation and sterol-responsive transcriptional regulation, which could significantly increase the yield of monoterpene (limonene or geraniol) production in engineered yeast strains [87].

Similarly, many protein down-regulation strategies have been already applied to Erg7p. Broker et al. inhibited the expression of *the ERG7* gene by the copper-regulated promoter, which resulted in the accumulation of high specific 2,3-oxidosqualene titer (about 200 mg/g DCW) in the presence of 150 μM CuSO_4_. However, the bulk of the accumulated 2,3-oxidosqualene seemed to contribute less to improve the production of triterpenoids [88]. Similarly, Kirby et al presented a similar strategy that repression of *ERG7* gene was achieved by the methionine-repressible promoter. However, they only observed a 50% increase in the yield of β-amyrin [16]. Moreover, the TetR-TetO based gene regulation system was used to manipulate the transcriptional efficiency of *ERG7* gene, which only led to an about 10% increase in the yield of protopanaxadiol [89]. It appears that the down-regulation of the *ERG7* gene has a limited effect on improving triterpenoid production.

### Challenges for triterpenoids production in yeast

To date, most of the metabolic engineering approaches focus on elevating the carbon flux of the MVA pathway. However, there are fewer successful cases on the manipulation of Erg7, OSCs, P450s, and UGTs, which are still challenges that need to be overcome. These challenges also represent the potential for continued improvement for the heterologous production of triterpenoids in *S. cerevisiae*.

Although microbial synthesis is relatively easier than the chemical synthesis, there are still many uncharacterized synthetic pathways of triterpenoids. Due to the loss of intermediates, it is challenging to speculate an unknown synthetic pathway based on the limited intermediates and the end-products. Even though several triterpenoid biosynthetic pathways have been identified and characterized, we still lack appropriate enzymes expressed in yeast. For example, a monofunctional α-amyrin synthase has not yet been identified in plants, which causes the production of α-amyrin frequently with the accumulation of β-amyrin, lupeol, and other triterpenes. In our sturdy, CYP716A15 catalyzed the conversion of lupeol to betulinic acid, with the concomitant of betulin and betulinic aldehyde due to incomplete oxidation. This phenomenon seriously influences the yield of targeted products and causes a high-cost purification process. Besides, there is currently a lack of a chassis capable of producing high-level production of 2,3-oxidosqualene, which is simultaneously beneficial for triterpenoids production. In general, squalene accumulation is a representative indicator for the carbon flux of the MVA pathway. However, no 2,3-oxidosqualene producing yeast strain has been achieved through metabolic engineering, probably due to that there is not an effective strategy for down-regulation of *ERG7* gene.

Although ergosterol is essential for yeast cell viability, a decrease of ergosterol content concurrent with 2,3-oxidosqualene accumulation can be realized by the addition of U14266A (3ß-(2-dimethylaminoethoxy)-androst-5-en-17-one, an inhibitor of Erg7p) [95]. This result indicated that there is some space to repress the Erg7p activity in the condition of healthy cells. Nonetheless, Erg7p is not a single enzyme that has complicated interactions with other enzymes in the sterol pathway, such as Erg6p, Erg11p, Erg27p [96]. For example, deletion of 3-keto sterol reductase (Erg27p) leads to a concomitant loss of lanosterol synthase (Erg7p) [97], it is probably a hurdle to manipulate the Erg7 for redirection the carbon flow to the triterpenoid biosynthesis rather than the native sterol pathway.

### Perspective

Identification of novel genes is a vital tool for understanding the triterpenoid synthetic pathway, especially as often more than one triterpenoid is synthesized and the enzyme classes (OSCs, P450s, UGTs) contribute to biochemical diversity. Besides, a transporter may exist in the plant cell, which can transport betulin and even betulinic acid to the bark of white bitch trees. A functional transporter probably reduces the metabolic burden, improves the production of triterpenoids and benefits for the extraction process in yeast. With the development of synthetic biology, high-throughput sequencing is a potential method to dig novel genes for efficient production of triterpenoids.

Apart from the discovery of the novel gene, application of protein engineering on the existing Erg7, OSCs, P450s, and UGTs is also a promising strategy to facilitate the triterpenoid biosynthesis. With the development of protein engineering, many mature strategies have been applied to control protein activity. For example, several conditional degrons have been designed to control the abundance of targeted protein by adapting experimental conditions, which is a promising strategy to repress the Erg7 expression [98, 99]. Active-site mutagenesis of triterpenoid-related P450s might increase the catalytic efficiency and decrease the generation of by-products, which has been utilized to improve the selectivity and catalytic activity of the bacterial P450s [100].

Considering that OSCs and P450s are respectively localized in lipid droplets and ER membranes [101], the co-localization of these enzymes is probably beneficial for the yield of triterpenoids. For instance, relocation of the MVA pathway to the mitochondria could enhance the carbon flux which further improved the yield of sesquiterpene [102]. Moreover, compartmentalization of isobutanol biosynthetic pathway into the mitochondria was also able to improve the isobutanol yields by 260% in *S. cerevisiae* [103].

In conclusion, *S. cerevisiae* is an excellent alternative for biotechnological production of valuable triterpenoids and identification of putative triterpenoid-related enzymes. Metabolic engineering and protein engineering allow further modification of yeast to efficiently produce targeted triterpenoids, which paves the way for the industrial production in place of extraction from plant sources.

## Conflicts of interest

The authors declare no conflicts of interests

## Acknowledgments

This research was supported by the National Natural Science Foundation of China (grant No. 21676026), and the National Key R&D Program of China (grant No. 2017YFD0201400).

